# Successful delivery of CRISPR-Cas9 with a baculovirus vector for insect brain targets

**DOI:** 10.1101/2024.11.26.624635

**Authors:** Büşra Elif Kıvrak Doğan, Saleh Ghanem, Michael Goblirsch, Farzana Nazneen, Robert L. Broadrup, Annie Connolly-Sporing, Doğa Tuncer, Christopher Mayack

## Abstract

CRISPR-Cas9 (clustered, regularly interspaced, short palindromic repeats with CRISPR-associated protein 9) is a powerful, versatile, and cost-effective molecular tool that can be used for genetic engineering purposes and beyond^1^ and is especially suited for non-model organisms^2^. Effective delivery of this system, however, remains a challenge for *in vivo* genetic manipulation of specific tissues^3^, particularly the brain^4^, and in adult indivuduals^5,6^. We designed a new CRISPR-Cas9 plasmid that was inserted into a baculovirus vector to knockdown the *octopamine beta subtype 2 receptor* (*AmOctβ2R*), a transmembrane protein found in the mushroom body neurons of the honey bee (*Apis mellifera*) brain, to determine if octopamine plays a role in appetite regulation. We first confirmed that gene editing of *AmOctβ2R* is possible with Sanger sequencing. We then demonstrated expression of the CRISPR-Cas9 system with the baculovirus vector *in vitro* using live cell imaging, flow cytometry analysis, and *in vivo* using confocal imaging, showing widespread expression in the cells and throughout the honey bee brain, three days post treatment. There was also *in vitro* and *in vivo* knockdown of *AmOctβ2R* three days post-infection, that corresponded with appetite suppression in starved forager bees. Our findings suggest that we successfully delivered the CRISPR-Cas9 system and knocked down *AmOctB2R* in neuronal cells of the honey bee brain that were previously inaccessible due to the blood brain barrier and lack of infectivity of lentivirus vectors^7^. The newly characterized *AmOctβ2R*^8^ can now be assigned a functional role and other targets for gene editing are now possible using this CRISPR-Cas9 system.

**Graphical Abstract:** 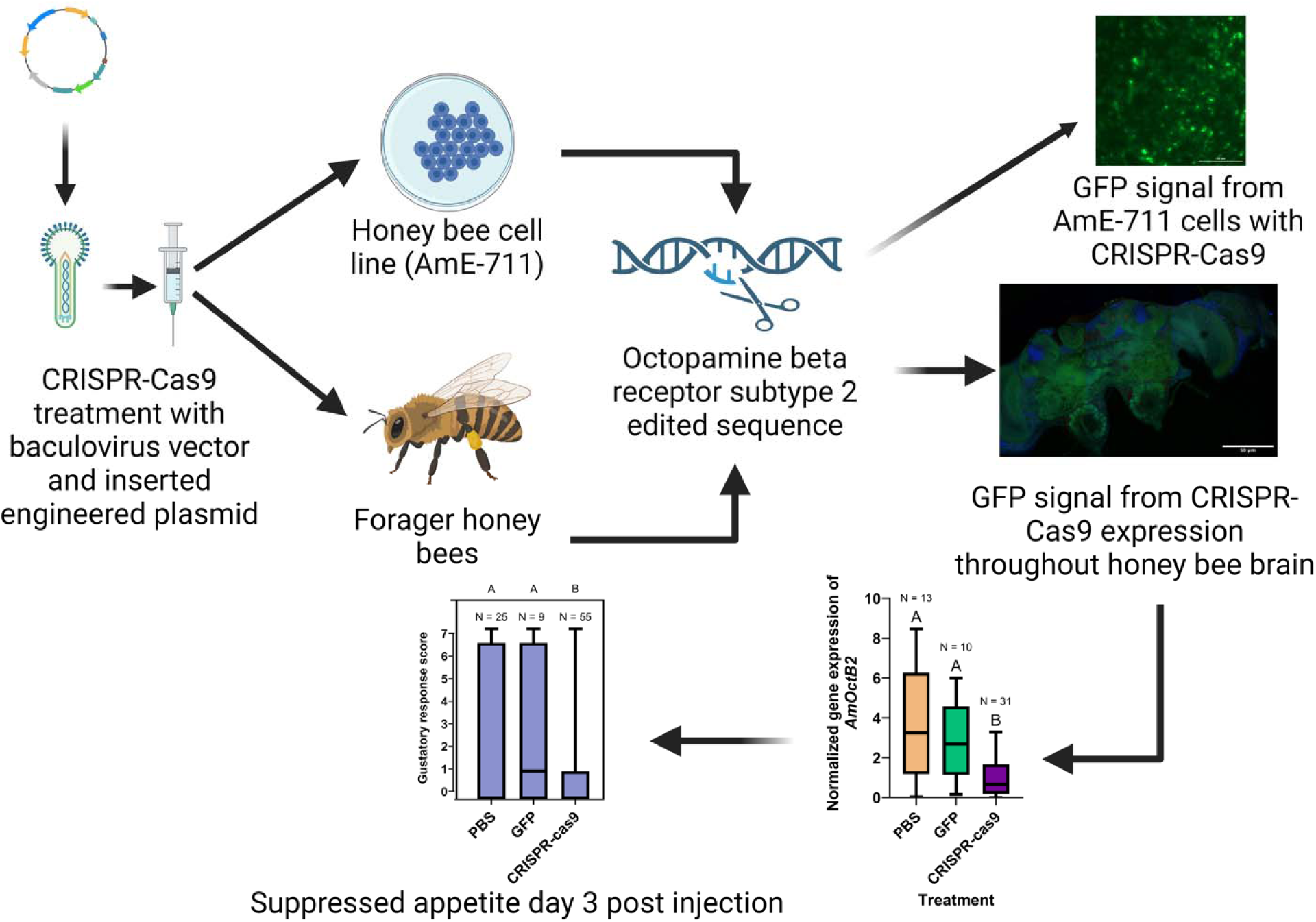

## Results

CRISPR-Cas9 (clustered, regularly interspaced, short palindromic repeats with CRISPR-associated protein 9) is a powerful, cost-effective, genetic engineering tool that is amenable for genetic manipulation in non-model organisms^1,2^. However, there is limited success of delivering the CRISPR-Cas9 system *in vivo*, impeding its potential impact^3,5,6^. This is especially the cas for less accessible targets in the brain, where the blood brain barrier inhibits diffusion of the CRISPR-Cas9 system and insensitivity of certain cells, such as Kenyon cells, to lentivirus infection^4,7^.

Recent improvements in gene editing approaches in insects^9^ have revealed the function of genes in life history^10^ and identification of olfactory receptors in neurons in transgenic individuals^11^. Moreover, using CRISPR-Cas9 for creating honey bee transgenic lines has remained challenging because they cannot be propagated in the laboratory^12^. Injection of genome editing materials into queen bee ovaries is difficult to achieve as honey bees do not have single germline stem cell like *Drosophila*. Instead, their germline progenitors are found in clusters, which would lead to a variety of patterns when the cells divide^13^. Despite these limitations, the honey bee, with its simple, yet true brain composed of functionally specialized tissues, has emerged as an ideal model organism for studying neurological functions that underlie complex social behaviors^14^, navigation^15^, olfactory processing^16^, color vision^17^, cognition^18^, and learning and memory^19^. Therefore, an effective gene delivery technique for the adult honey bee brain would provide a valuable approach in which gene editing could be achieved using the CRISPR-Cas9 system.

Here, we chose the baculovirus as the viral vector over the common lentivirus that is typically used with mammalian organisms because it has a higher infectivity rate for insect cells and presents lower risk to human health to deliver the CRISPR-Cas9 system to knockdown the *octopamine beta receptor subtype 2* (*AmOctβ2R*) neuronal transmembrane octopamine signaling protein to determine its role in appetite regulation^20,21^. We first confirmed that various guide RNAs (Table S1), when injected into adult bees with the CRISPR-Cas9 complex, could successfully edit all four *octopamine beta receptor subtypes 1-4* (Table S3). We then delivered the CRISPR-Cas9 system into the brains of honey bee foragers using a baculovirus vector with an integrated plasmid named as pBV-rev(U6-3>gRNA)-Am-actin5c>DmCas9:T2A:EGFP:Puro) (VectorBuilder, Chicago, IL, USA: VB210408-1080vfc) that was designed to mutate and lower the expression of *AmOctβ2R* (Fig. S1). We performed ocellar tract injections using the vector and exhibited that this receptor plays a role in gustatory processing by demonstrating appetite suppression in starved individuals starting one day after injection (Chi-square goodness of fit day 1: df = 2; χ^2^ = 12.6; P = 0.0018) and a maximum effect observed three days after injection (Chi-square goodness of fit day 3: df = 2; χ^2^ = 27.47; P < 0.0001; Fig. 1). We also validated successful knockdown of *AmOctβ2R in vivo* by demonstrating significantly lower gene expression in CRISPR-Cas9 treated bees, three days post injection (Kruskal-Wallis Test: χ^2^_2,53_ = 13.47, P = 0.0012; Fig. 2). Successful injection of the CRISPR-Cas9 baculovirus system was further confirmed by confocal microscopy, where enhanced green fluorescent protein (eGFP) signaling was substantially higher throughout the honey bee brain. This imaging also provided evidence of transduction and expression of the CRISPR-Cas9 system in Kenyon cells of the honey bee brain, which is where the *AmOctβ2R* transmembrane signaling protein is located (Fig. 3). Treatment of the *Apis mellifera* cell line,AmE-711, with the CRISPR-Cas9 complex yielded knockdown of *AmOctβ2R* three and four daysafter infection relative to the PBS and eGFP controls (Kruskal-

**Figure 1.**
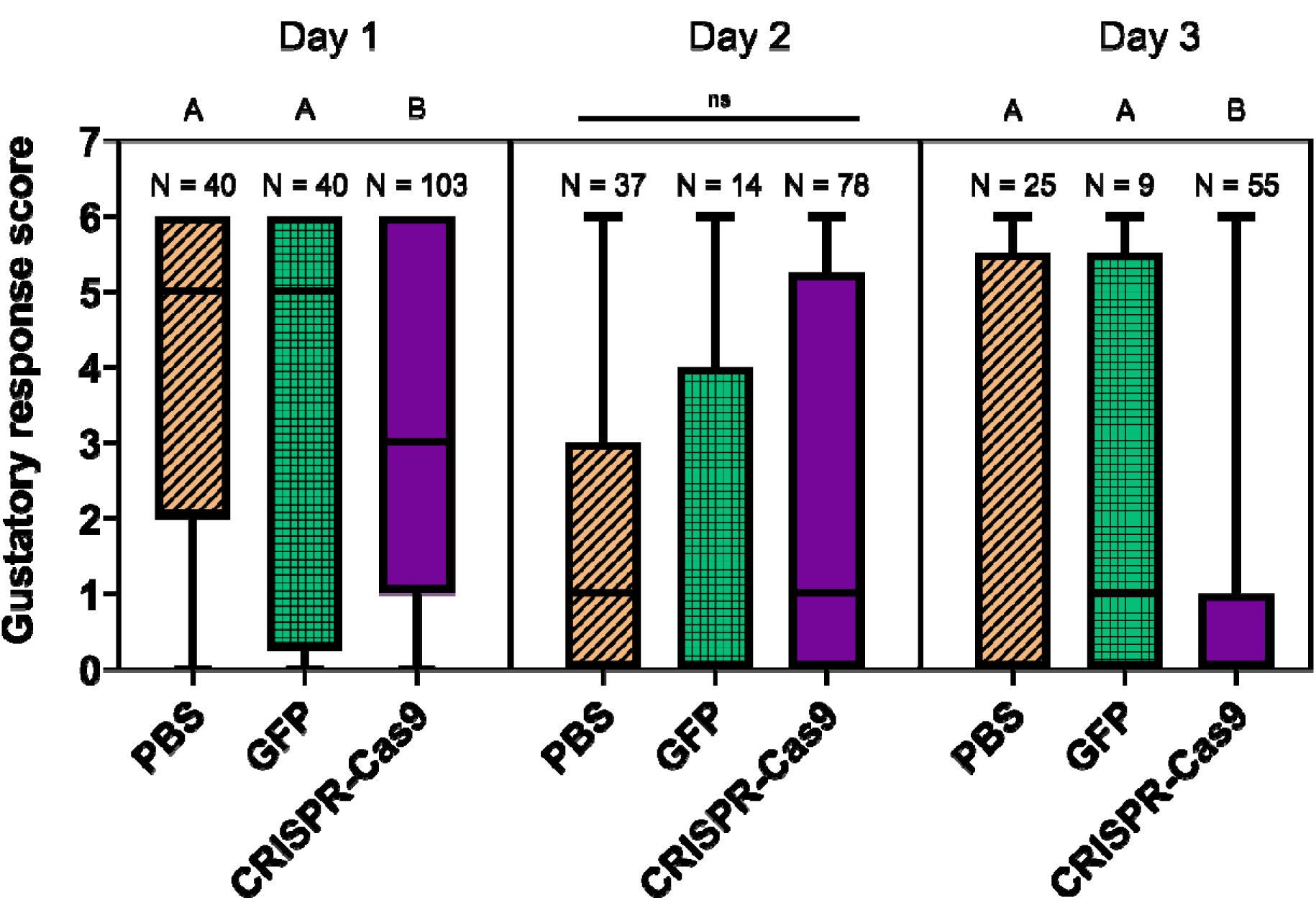
Gustatory response scores of honey bees (*Apis mellifera*) 1- 2-, and 3-days post ocellar tract injection. Treatments were 1x phosphate buffered saline (PBS), enhanced green fluorescent protein reporter in 1x PBS (eGFP), and baculovirus containing the CRISPR-Cas9 *AmOctβ2R* knockdown plasmid (CRISPR-Cas9) in 1x PBS. Each box represents the inter quartile range, the middle horizontal line represents the median, and the whiskers represent the minimum and maximum range. The letters above the boxes represent significant differences at the P < 0.05 level. Sample sizes are listed above each bar for each treatment group.

**Figure 2.**
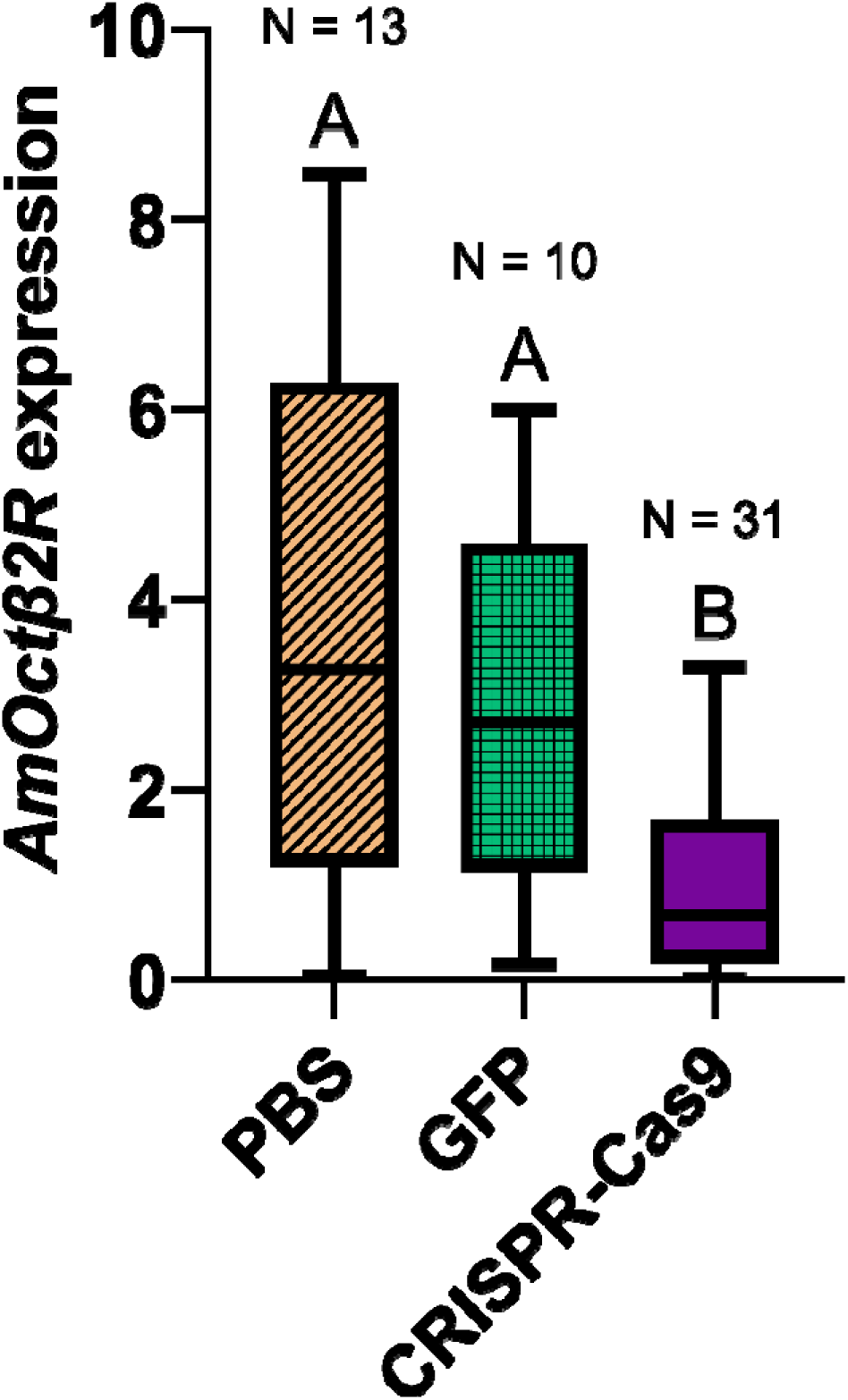
Normalized expression of *octopamine beta receptor subtype 2* (*AmOctβ2R*) from the honey bee (*Apis mellifera*) head three days post ocellar tract injection. Treatments were 1x phosphate buffered saline (PBS), enhanced green fluorescent protein reporter in 1x PBS (eGFP), and baculovirus containing the CRISPR-Cas9 *AmOctβ2R* knockdown plasmid (CRISPR-Cas9) in 1x PBS. Boxes represent the interquartile range with the middle horizontal line representing the median; whiskers mark the minimum and maximum values. Letters above the boxes represent significant differences between the treatment groups, α = 0.05. Sample sizes for each group are indicated above the letters.

**Figure 3.**
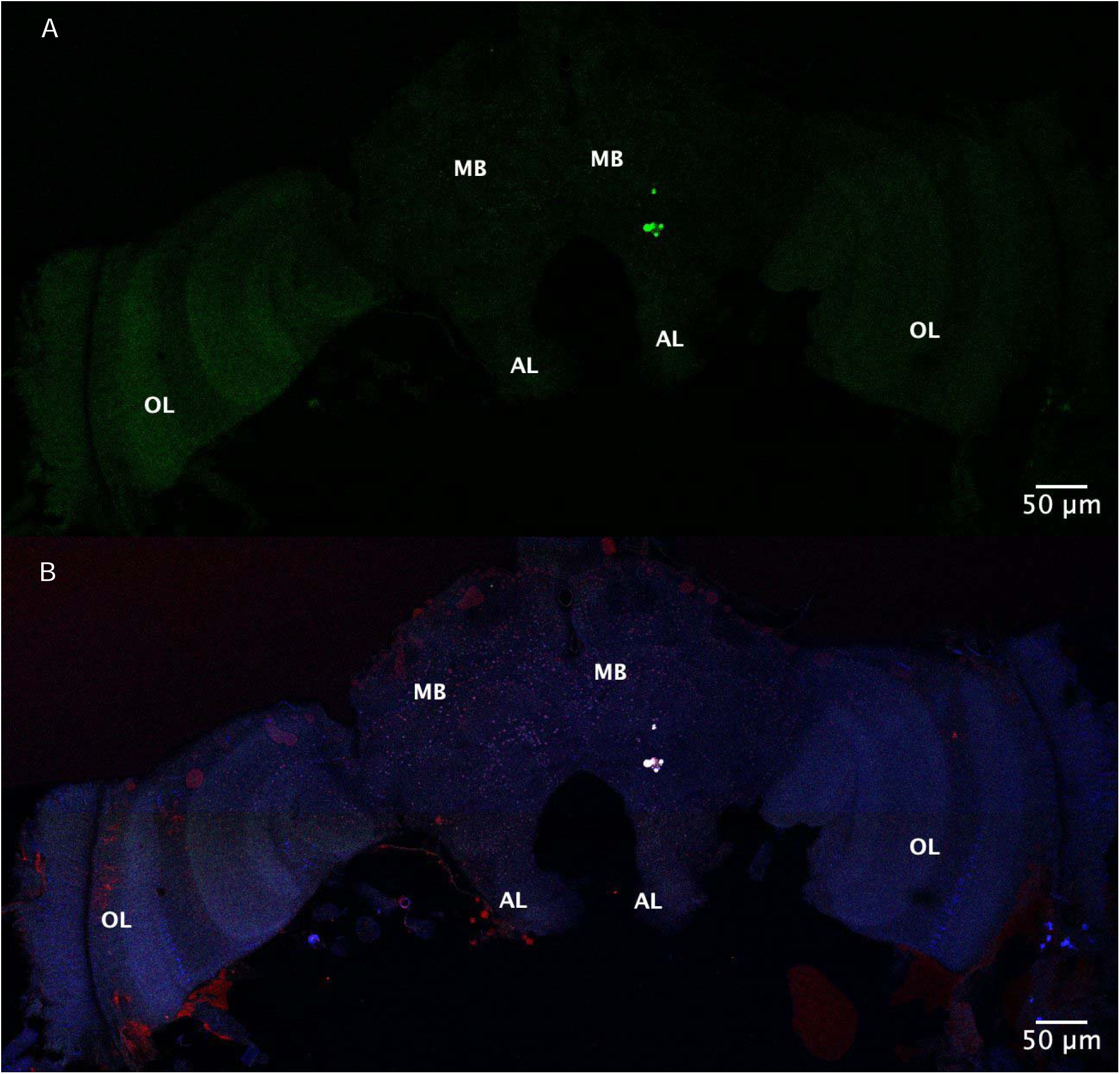
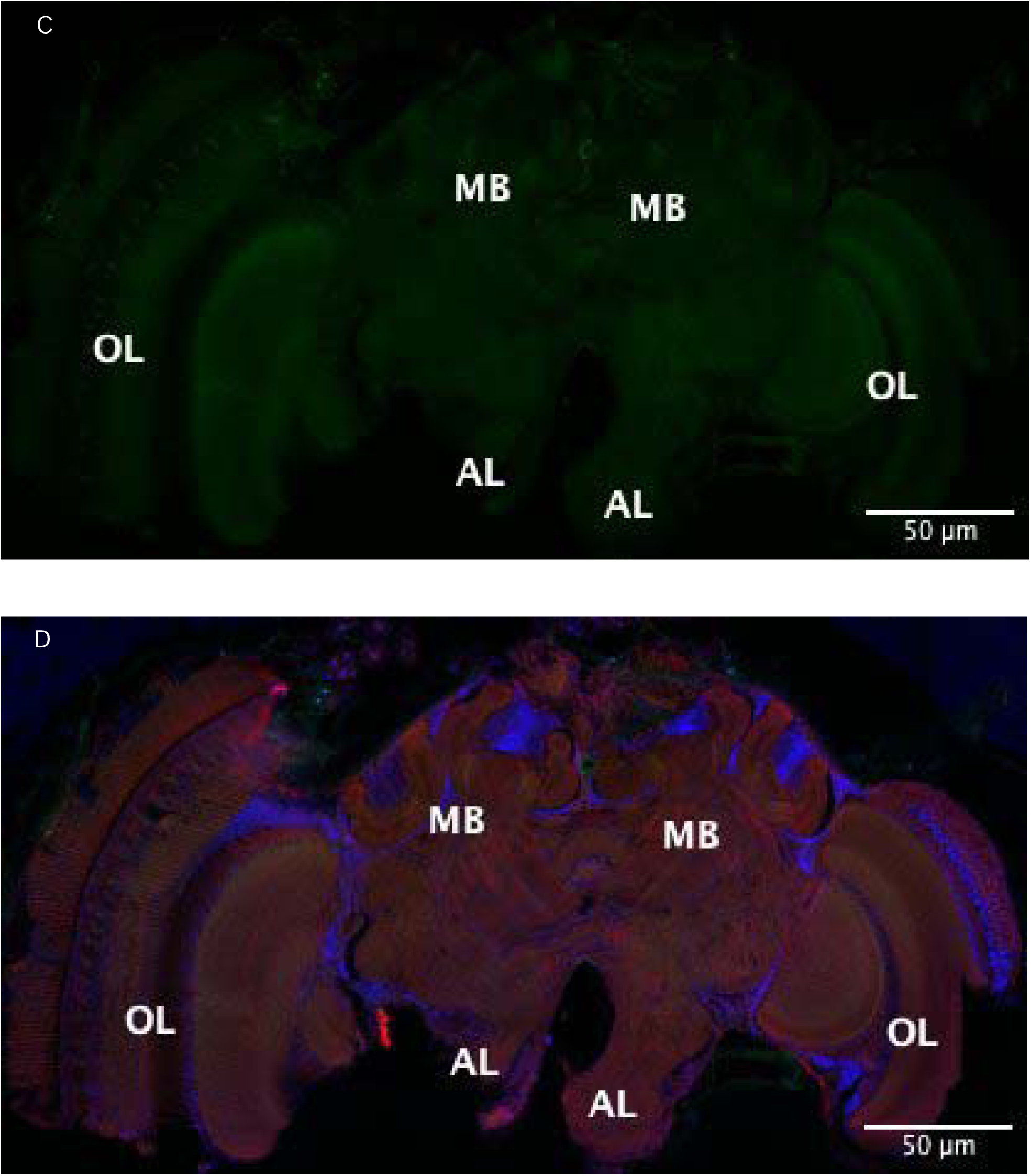
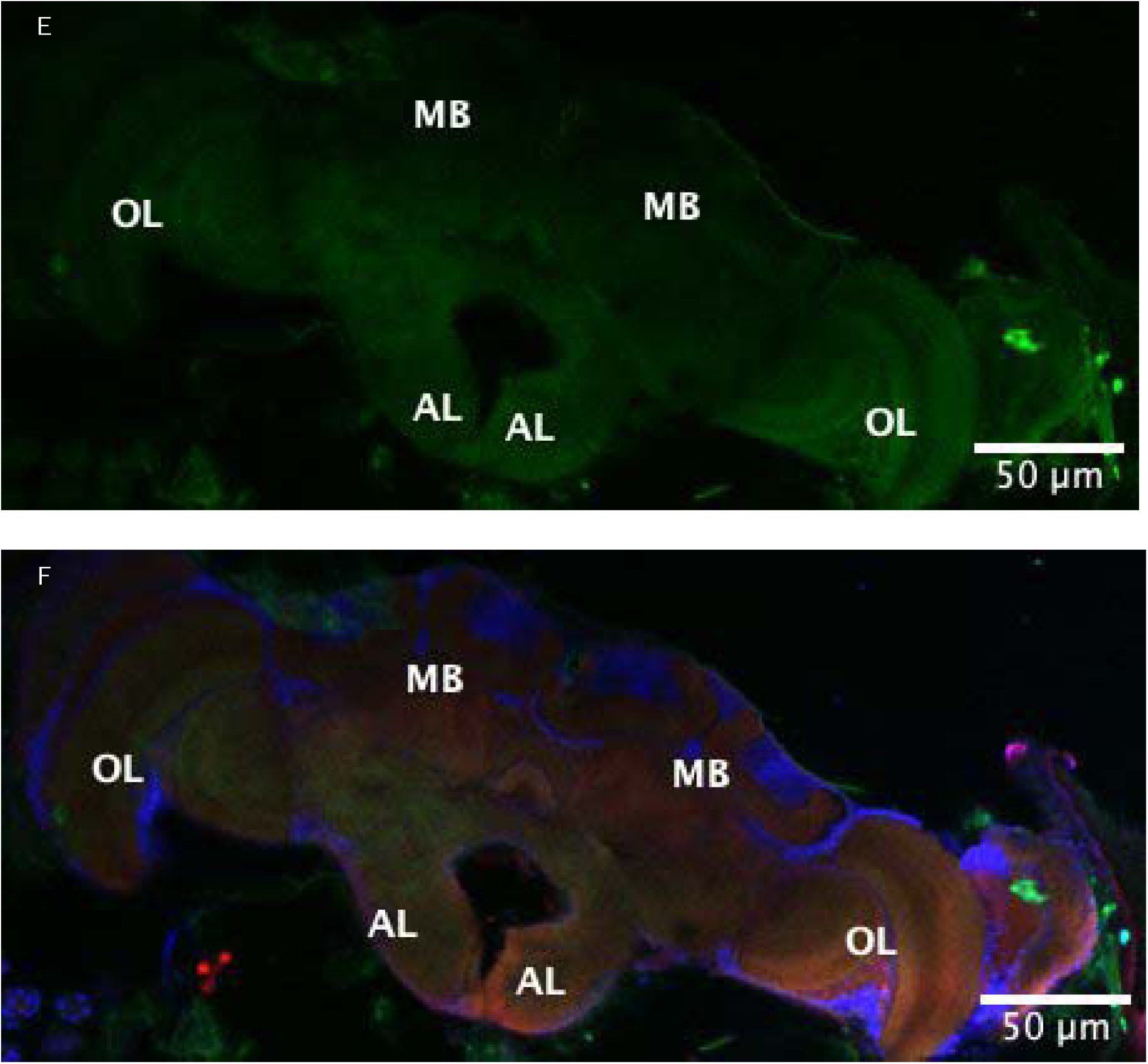
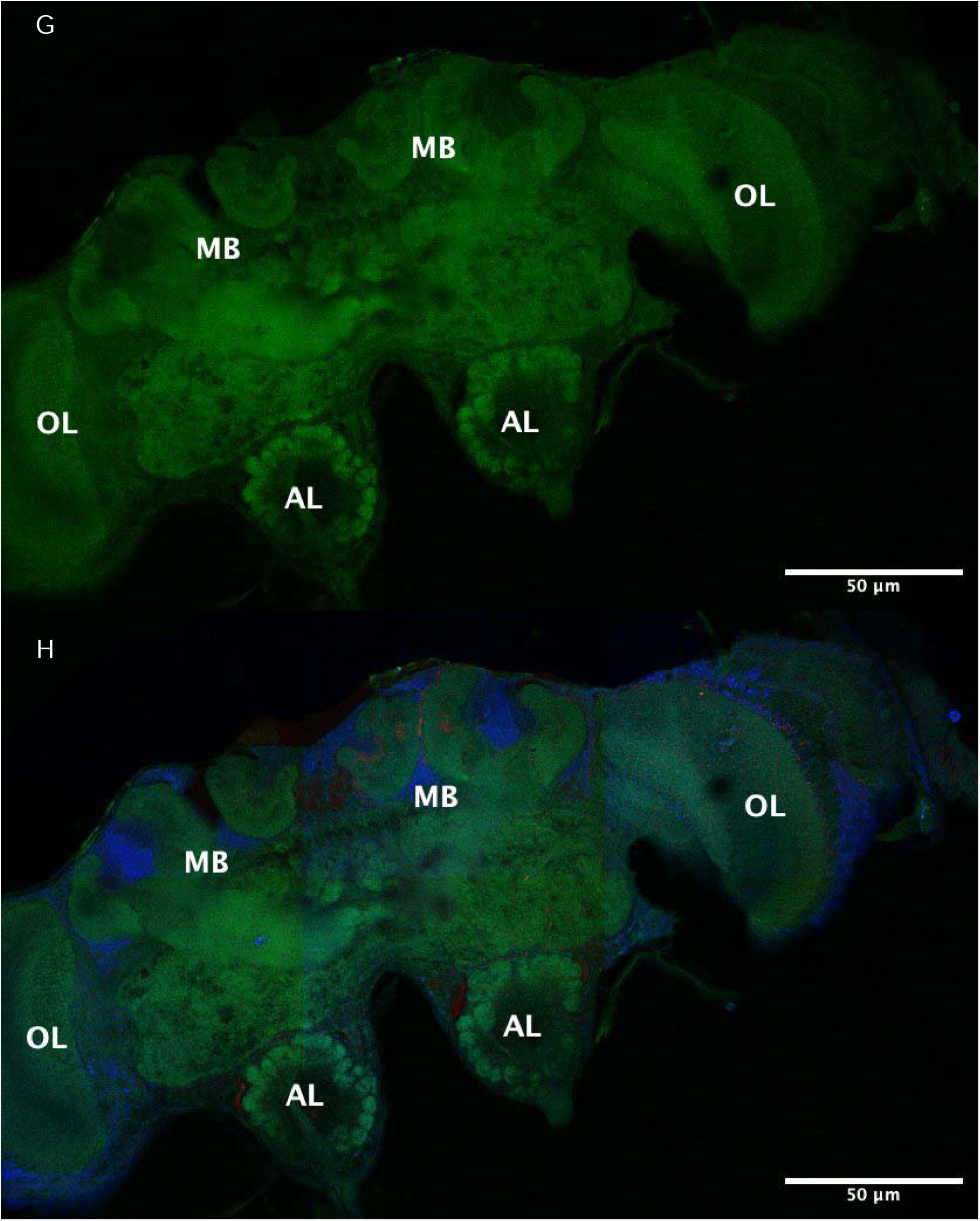
Comparison of eGFP signal in CRISPR-Cas9 injected and control honey bee brains. Images of noninjected whole honey bee brains for enhanced green fluorescent protein expression (eGFP) signal only (green) (A) and then for all channels (B), where DAPI for cell nuclei staining is shown in blue and phalloidin for actin staining is shown in red. Phosphate buffer solution 1x (1x PBS) injected three days post injection for the Green Fluorescent Protein expression (GFP) signal only (green) and then for all channels (C and D, respectively). CRISPR-Cas9 injected whole honey bee brains two days post injection for GFP (E) and all channels (F), respectively. CRISPR-Cas9 injected whole honey bee brains three days post injection for GFP (G) and all channels (H), respectively. The antennal lobe (AL), mushroom body (MB), and optical lobe (OL) regions of the whole brain are indicated for spatial reference. The images are showing the whole brain with a slice thickness of 50 µm. The voltage gain for all images was set to 650 dB.

Wallis Test: day 3 after infection: χ^2^ = 19.64, P = 0.0015; day 4 after infection: χ^2^ = 17.17, P = 0.0042), all treatment groups, especially the PBS control group, also demonstrated a significant increase in *AmOctβ2R* gene expression over time (Kruskal-Wallis Test: Time: χ^2^ = 73.30, P < 0.0001) (Fig. S2). On day 3 post-infection we confirmed a dose dependent increase in eGFP signal for CRISPR-Cas9 treated cells as indicated from flow cytometry analysis (Fig 4.).

**Figure 4.**
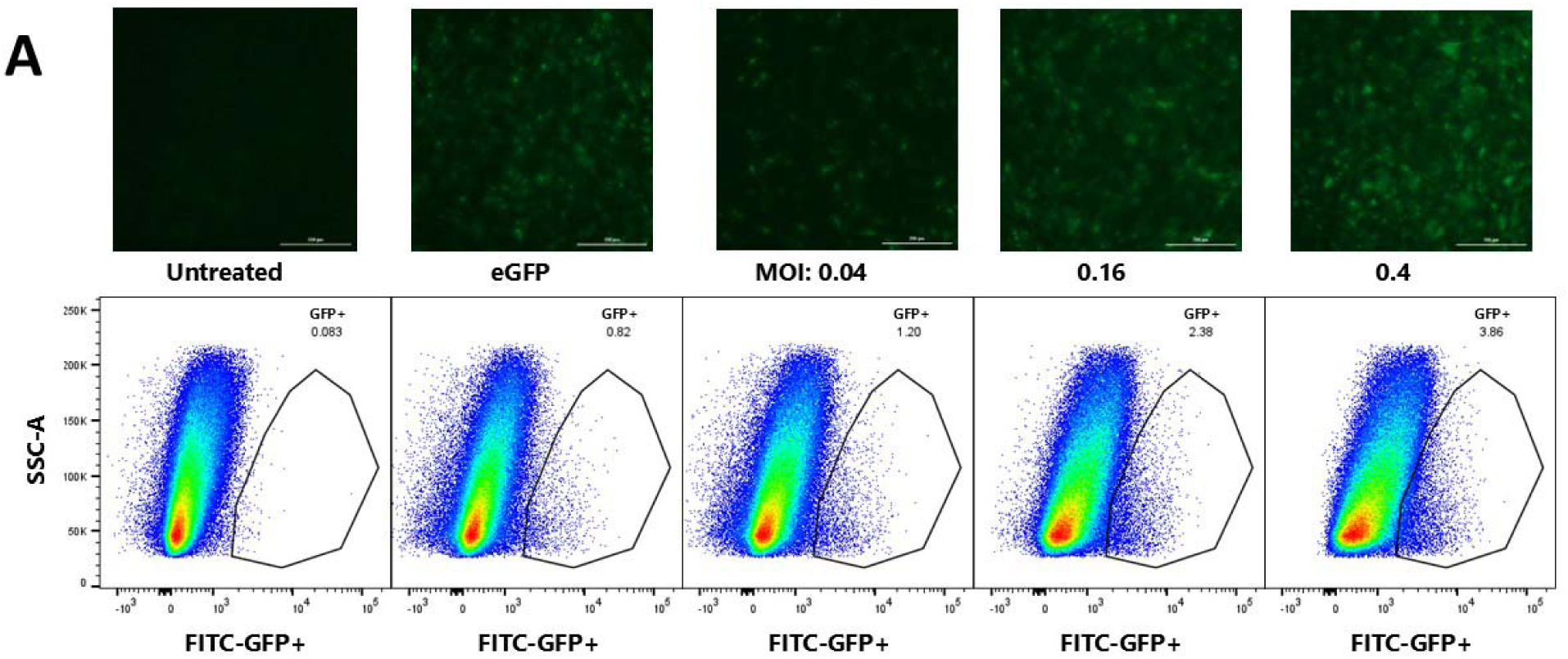
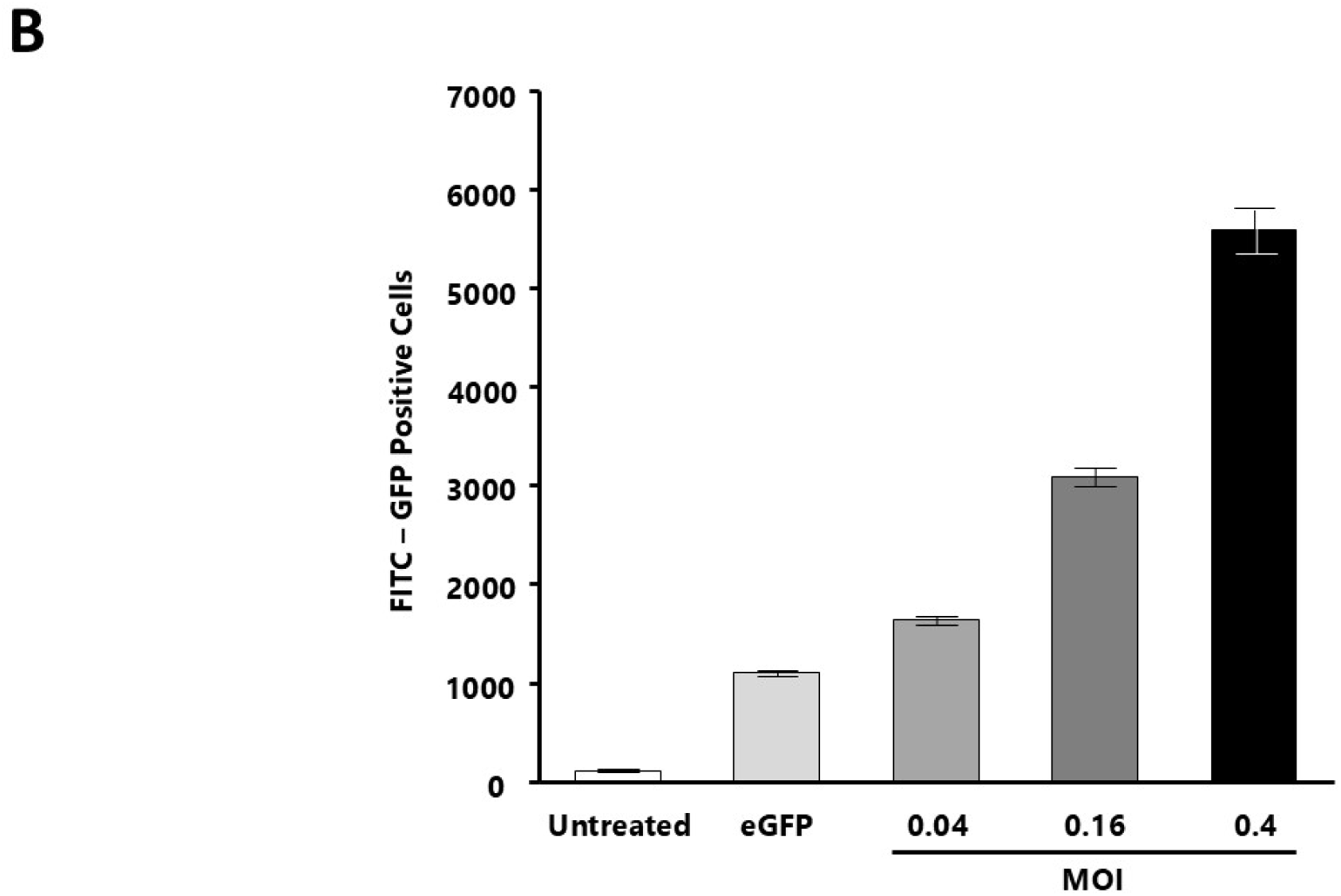
AmE-711 honey bee cells were characterized for expression of enhanced green fluorescent protein (eGFP) by live cell imaging, shown above each label, and flow cytometry, 72 hours post treatment (A). The percent eGFP positive cells within the gated population is shown in the upper right of a representative panel from each treatment group. Bars represent the mean number of GFP positive cells ± SD of three wells per treatment group (B).

## Discussion

Gene editing approaches for targets in the honey bee brain have been largely unsuccessful. Electroporation has resulted in high mortality levels^22^, whereas the lentivirus vector that is typically used for vectoring CRISPR-Cas9 in mammalian cells, was unable to infect the neurons of the honey bee brain^7^. In general, viable vectoring systems have been sparsely tested in honey bees. Most previous work for gene editing of the honey bee using CRISPR-Cas9 has been carried out by direct injection of embryos^23^. However, this is less amenable when targeting genes that might lead to a loss of survival over time^24^ or underlie behaviors that are more transient in adults^25^. In addition, direct injection of adult honey bee brain using RNAi gene silencing has resulted in limited penetration of the gene delivery agent and low efficiency levels^26^. Therefore, there is need for improving the CRISPR-Cas9 system for valuable *in vivo* gene editing in adult insects, especially for targets in the honey bee brain.

Roughly 70% of the samples from the different guide RNAs across the *AmOctβR 1-4* targets produced substantial mutations of the intended target sites. We detected CRISPR-Cas9 expression in Kenyon cells of the mushroom bodies^7^, as well as other regions of the honey bee brain, which confirms that the CRISPR-Cas9 system is able to reach the *AmOctβ2R* target, which is a transmembrane protein found in honey bee neuronal cells^8^. We then confirmed expression of the CRISPR-Cas9 system within the honey bee cells by observing a dose-dependent response based on the multiplicity of infection (MOI) of baculovirus vector administered with live cell imaging, but also quantitatively with flow cytometry. We also found significant *in vitro* and *in vivo* knockdown of *AmOctβ2R* three days after injection, which corresponded with the largest amount of appetite suppression in starved forager bees.

Taken together, our findings suggest successful delivery of the CRISPR-Cas9 system and knockdown of *AmOctβ2R* in neuronal cells of the honey bee brain, a tissue that was previously found to be inaccessible to gene editing due to the blood brain barrier and insensitive to other viral vectors^7,27^. By effectively knocking down the *AmOctβ2R*, we were able to assign a new function related to appetite regulation to this relatively recently characterized honey bee octopamine receptor subtype^8^. Previously, a baculovirus- GFP expression system was reported to have limited distribution around the injection site in honey bee queen pupae, and did not reach the ovaries, which was the intended tissue target^28^. However, our confocal images of CRISPR- Cas9 injected honey bee brains displayed widespread eGFP signaling two and three days after injection, suggesting that the baculovirus vector was very efficient in infecting cells throughout the brain. The spread of the CRISPR-Cas9 treatment is critical because *AmOctβ2R* is thought to be present throughout the brain^29^. This finding is contrary to the lentivirus vector, which takes longer to express the CRISPR-Cas9 system and could not infect Kenyon cells of the honey bee brain^7^. Yet, these cells would be most desirable to manipulate using CRISPR-Cas9 because they play many functional roles in synaptic transmission and are found in the mushroom body region, which is responsible for processing sensory information^30^. Baculovirus vectors have efficiently transduced mosquito cells, without viral propagation, resulting in a high level of gene expression two days post injection when coupled with suitable promoters^31^. On the other hand, gene delivery using the adeno associated virus (AAV) vector has lower transduction efficiency, and may take two to seven days after injection to be expressed, resulting in a less practical option for CRISPR-Cas9 delivery^32,33^. Given how quickly and efficiently the baculovirus vector spread throughout the honey bee brain, our results suggest that a lower dose may have the same knockdown efficacy. Supporting this notion, we did not observe dose dependent knockdown in the honey bee cells when infecting them with increasing MOI of the baculovirus.

Gene editing approaches for honey bees have previously been conducted on queens, eggs, fat body, and sperm with the goal of creating transgenic lines that would eliminate the primary challenge of maintaining mutant organisms in honey bee genetic studies^28,34,35^. However, previous attempts of gene editing the honey bee germ line has not had a high level of success. Artificial insemination of sperm transduced with linear DNA resulted in negative impacts on metamorphosis of the next generation of bees and there was limited transfection success. In addition, the introduced DNA was not integrated into the genome of the next generation, as a result, knockout lines of honey bees are not available^35^. The first CRISPR-Cas9 application on honey bee embryos was successful^36^, but these individuals had to be raised indoors due to biosecurity regulations^37^. Therefore, the current method for delivering CRISPR-Cas9 has been injecting larvae or pupae for loss of function studies. However, injection at these life stages resulted in high mortality before these individuals could reach adulthood^38^.

Octopamine release is known to be a part of the general stress response in insects^39^, which may also explain why we observed the increased gene expression of *AmOctβ2R* in the PBS injected control bees *in vivo* three days post injection. From our *in vitro* assays, we found a general increase in *AmOctβ2R* gene expression over time. This is likely because the cells became stressed in the 24-well plate conditions, as we noticed cell mortality, especially in the PBS treated cells. Based on these findings, for future work, we would recommend finding another solvent besides PBS for the baculovirus vector, or at least diluting the PBS, before carrying out future experiments on insects.

There have been previous attempts for reaching gene targets in the honey bee brain *in vivo*. For example, eGFP delivery with Am-actin5c, elp2l, Am-hsp83 and Am-hsp70 promoters was achieved through electroporation, but again, there was around 50% mortality after treatment and the transfected region was limited to the edges of the brain, suggesting limited penetration.^22,40^. Therefore, there is large potential for use of the baculovirus vectors to deliver the CRISPR-Cas9 system. Here we used it to unveil a pathway involved with appetite regulation in the forager honey bee. Previous work has demonstrated a correlation between starvation, hemolymph trehalose levels, octopamine levels in the brain, and appetite regulation of honey bee foragers^41,42^ and this newly recognized regulatory pathway appears to be independent of the glucose-insulin nutrient signaling pathway of honey bees, that interestingly, in contrast, is commonly found in vertebrate appetite regulatory systems^43^. We demonstrated a cause-and-effect relationship between octopamine signaling and appetite regulation through delivery of a baculovirus- facilitated CRISPR-Cas9 system into the honey bee brain. Furthermore, we provided proof of concept for this gene editing method to become a practical neurobiological tool, which could be suitable for uncovering other neurobiological mechanisms, for a range of well-studied honey bee behaviors, such as complex social behaviors. More recently *AmOctβ2R* has been shown to play a role in honey bee thermoregulation and is expressed both in the muscle and brain^44^. Consequently, our CRISPR-Cas9 system could be used to unveil further functional roles this receptor plays for honey bee as it has pleiotropic effects.

The baculovirus vector is particularly advantageous due to its large cargo capacity (up to 38 kilobases (kb)), and as such, it holds potential to be used with multigene delivery applications^45^. Currently, this vector is being engineered for use on mammalian cells because it is replication-incompetent and does not integrate into the host genome^45^. In addition, helper viruses are not needed, therefore, baculovirus vectors are relatively simple and easier to construct. Furthermore, baculovirus transduction levels can be up to 100%, with a very high expression level, and can be noncytotoxic, even at high MOI levels^46^. The advanced genome engineering toolbox, which includes constructs with “dead” Cas9, or dCas9, for transcriptomics could also be used^47^ for inducible genome editing systems which enables the turning on and off gene expression of specific gene targets. Therefore, dCas9 could be used under different chemical conditions in future studies to the examine role of several genes involved in signaling pathways^48^. Here we demonstrate that the use of the baculovirus vector for insect brain targets has the potential to make the powerful genetic engineering capabilities of the CRISPR-Cas9 system a practical reality, and our system is especially suitable for insect model organisms but could also be applied to a variety of other organisms.

## Materials and Methods

### 1. Honey bee collection, harnessing, and injections

From three different source colonies, we collected only bees with full pollen baskets to ensure that they were forager aged. We placed wire mesh at the entrance of three different hives and individually captured 50 pollen foragers at a time in 20 ml glass scintillation vials (Sigma-Aldrich, St. Louis, Missouri, United States). We placed the vials on ice to immobilize the bees while they were being transported to the laboratory. Next, we individually harnessed up to 50 bees at a time by placing them in cut plastic drinking straws which were approximately 5 cm in length. We stabilized the bees in the straws with a 1 mm width strip of duct tape that was placed in between their thorax and head. After 30 min of acclimation, we fed the harnessed bees with 50% sucrose solution *ad libitum* to standardize their hunger levels^49^.

In a given trial, honey bees were randomly divided into three groups of roughly 15 bees. The heads were stabilized with a collar made of dental wax and the lens of the middle ocellus was removed using a micro scalpel (Feather, Osaka, Japan) with a 30° blade. The first negative control group underwent a medial ocellar tract injection using a 10 μl Hamilton syringe (Hamilton, USA) placed in a micromanipulator (WPI, Germany) to inject 1 µl of 1x Phosphate-buffered saline (PBS) into the brain^50^, the second negative control group was injected with 1 µl of 1 x 10^7^ PFU/ml baculovirus vector with eGFP reporter, and the third treatment group was injected with 1 μl of 1 x 10^7^ PFU/ml baculovirus vector with the CRISPR-Cas9 plasmid insert. All vectors were dissolved in PBS. After injections, the harnessed bees were once again fed *ad libitum* with 50% sucrose solution. Then, they were placed inside an incubator at 25°C and 70% relative humidity when not being used for the experiment.

### 2. Proboscis Extension Response (PER) assay to measure appetite levels

We measured the appetite levels of the experimental bees daily for 3 days using the Proboscis Extension Response (PER) assay^51^ . Prior to the assay on each day, bees were starved for 18 h. We performed the PER assay by touching a droplet of sucrose solution to the bee’s antenna in ascending order of 0.1%, 0.3%, 1%, 3%, 10%, and 30% concentrations. In between each concentration, a droplet of water was touched to the antenna of the honey bee to prevent desensitization. Scoring of the PER was binary, a score of 1 was recorded if the bee fully extended its proboscis in response to the droplet of sucrose solution and a score of 0 was recorded if the bee showed no full proboscis extension. A Gustatory Response Score was calculated from the PER assay by summing scores across all concentrations with a score of 6 as the maximum possible value^42^. After each PER assay, we fed the bees *ad libitum* with 50% sucrose solution and then also 6 h later to maintain 18 h of starvation until the next day. After the third day, the bees were flash frozen in liquid nitrogen and their heads were removed and stored at -80°C until further qPCR analysis.

### 3. In vitro assay

AmE-711 honey bee cells were used with permission from the University of Minnesota through an agreement with the USDA. Cells were seeded at an approximate density of 2.5 x 10^5^ per well of a 24-well plate, with one plate used for each timepoint. Cells were allowed to recover from trypsinization and acclimate to the multi-well plates for two days prior to exposure to the treatment media. On day three after seeding, the culture media was removed from the wells and replaced with the corresponding treatment media, which consisted of: 1) Schneider’s Insect Medium alone as an uninfected/untreated control; 2) 1x PBS alone as a vehicle control; 3) baculovirus vector containing enhanced green fluorescent protein (EGFP) reporter construct as an expression control; and 4) baculovirus vector containing the CRISPR-Cas9 AmOct*β*2R construct with eGFP reporter plasmid insert at three different MOIs. The MOIs for the baculovirus AmOct*β*2R vector were approximately 0.04, 0.16, and 0.4. All treatments were diluted in Schneider’s Insect Medium, without fetal bovine serum, except for the PBS control. Cells were incubated in the treatment media for 24, 48, 72, or 96 h at 32 in a non-humidified incubator. An additional plate was used to establish baseline expression levels and consisted of cells exposed to the treatment media for 15 min (i.e., 0-h timepoint). At the set timepoints, the treatment media were removed and the cells were lysed in ice cold Trizol reagent (Thermo Fisher Scientific, Waltham, MA, USA). Total RNA was extracted from the lysates according to the Trizol manufacturer’s protocol except for inclusion of an additional 0.3M sodium acetate purification step. Total RNA was frozen at -80 and shipped on dry ice to the USDA facility at Davis, CA for further analysis.

### 4. qPCR gene expression analysis of octopamine beta subtype 2 receptor (AmOct*β*2R)

For the adult bees, an EcoPURE Total RNA kit (EcoTECH Biotechnology, Turkey) was used for RNA isolation by following the manufacturer’s instructions. Briefly, bee head samples were placed in a grinding specific, Safe-Lock 2 mL Eppendorf tube (Hamburg, Germany). Then, 300 μl of ECOPURE Lysis/Binding Buffer was mixed with 10% (3 μl) *β*-mercaptoethanol to inhibit the RNase activity in the samples. This mixture was added to each tube. A stainless-steel grinding bead was added to each centrifuge tube to be used in a tissue grinder (Qiagen, TissueLyser II). The mixture was then centrifuged for 10 min at 12,300 g to pellet the debris. The supernatant was transferred to a 1.5 ml microcentrifuge tube containing an equal amount of absolute ethanol (96-100%) and vortexed for 10 s. The mixture was transferred to the spin column and centrifuged at 12,300 g for 30 s at room temperature (RT). Flow through was discarded, and EURX DNAse (Gdańsk, Poland) treatment was followed using the manufacturer’s instructions.

Ten μl of DNAse treatment was added to the silica gel column for each sample. This was followed with a 30 min incubation period at 37° C. A total of 300 μl of EcoPURE wash buffer 1 was added to the column and centrifuged at 12,300 g for 30 s at RT. The flow through was discarded and 500 μl EcoPURE wash buffer 2 was added to the column, which was centrifuged at 12,300 g for 2 min at RT. The flow through was discarded and 200 μl of EcoPURE wash buffer 2 was added to the column. To remove any residuals, the buffer was centrifuged at 12,300 g for 2 min at RT. The column was then transferred to a sterile 1.5 ml microcentrifuge tube, and 35 μl of EcoPURE elution buffer was added to the column. The solution was centrifuged at 12,300 g for 2 min at RT. Flow through was retained and the previous step was repeated. The purity and concentration of the extracted RNA was measured for each sample using a Nanodrop spectrophotometer (Hampton, USA) and gel electrophoresis. The eluted RNA was then stored at -80 °C until further analysis.

For both the adult bees and cell samples, the cDNA was synthesized using a OneScript Plus cDNA Synthesis Kit (Abm, Canada), according to the manufacturer’s instructions. RNA was standardized to 35 ng/µl and a final volume of 15 µl before synthesizing the cDNA. For both the target gene, *AmOctβ2R*, and the reference gene, *Ribosomal protein 49* (*RP49*), 10 μl qPCR reactions were prepared consisting of 2.4 μl of nuclease free water, 0.3 μl of forward and reverse primers (SenteBioLab, Ankara, Turkey) (Table S1), 5 μl of Blastaq Green 2x master mix (Abm, Canada), and 2 μl of template cDNA. *RP49* expression was previously demonstrated to be stable across tissues and within the brain of the honey bee, *A. mellifera*^52^.

A Roche LightCycler 480 II thermocycler was used for the qPCR assay. Parameters were set to 45 quantification cycles (95, 30 sec ramp rate 2.2, 55 ramp rate 2.2, 30 s, 60 ramp rate 4.4, 1 min). We checked the specificity of the primer sets with a melt curve analysis (5 s, 95, 1 min, 60) and all samples were run in technical triplicates. *AmOctβ2R* expression was measured relative to *RP49* using the -ΔΔCT analysis method^39^.

### 5. Flow cytometry analysis for GFP expression

Cells were seeded at an approximate density of 2.0 x 10^6^ per well of three 6-well plates. Cells were allowed to recover from trypsinization and acclimate to the multi-well plates for two days prior to exposure to the treatment media. On day three after seeding, the culture media was removed from the wells and replaced with the corresponding treatment media and consisted of: 1) Schneider’s Insect Medium alone as an uninfected/untreated control; 2) 1x PBS alone as a vehicle control; 3) baculovirus vector containing eGFP reporter construct as an expression control; and 4) three different MOIs for the baculovirus vector containing the CRISPR-Cas9 AmOct*β*2R plasmid insert. At the end of a 72-hour exposure period, cells were collected from the multi-well plates using a cell scraper and samples placed on ice. The cells were washed twice with wash buffer. The cells were then fixed with 4% paraformaldehyde (PFA) by incubating for 15 min on ice in the dark. After washing once, the cells were resuspended in 300 µL staining buffer. Data were acquired using a LSRFortessa Cell Analyzer (BD Biosciences) and analyzed with FlowJo software (v10.9.0.).

### 6. Confocal brain imaging

#### 6.1 Sample preparation

Harnessed bees were placed head first on ice. We decapitated bee head and immediately placed it in 100 μl brain Ringer’s solution (130 mM NaCl, 5 mM KCl, 4 mM MgCl2, 5 mM CaCl2, 15 mM Hepes, 25 mM glucose, 160 mM sucrose, pH 7.2) on a dissection plate. Dissection of the brain was performed under a dissecting scope (Zeiss Stemi 305, Oberkochen, Germany) at 30x magnification to ensure that we did not lose any part of the brain tissue.

Dissected fresh brains were placed in a 1.5 ml microcentrifuge tube for fixation. The tube contained 1 ml of 4% PFA solution and was protected from UV light with aluminum foil. The brains were kept in 4% PFA overnight at 4° C. The brains then were kept in a 15% sucrose, PBS solution and then a 30% sucrose-PBS solution, respectively, until the brain sank to the bottom of the tube. The brains were rinsed with PBS for 10 min and then embedded in low melting point agarose (4%). Later, the brains were sliced (50 μm thickness) with a Leica 1,000s vibratome (location) at room temperature. These slices were kept in PBS at 4 °C until further analysis.

#### 6.2 Staining and confocal imaging of the honey bee brain

A staining solution was prepared with 0.4 U (units), equivalent to approximately 2 μl of phalloidin, and 1 μl of DAPI per slice. Phalloidin enables visualization of the actin filaments and DAPI binds to DNA, making the nuclei of the neurons visible. Slices were kept in staining solution for 20 min. Then, the brain slices were placed on a microscope slide and mounted with 20 μl of methyl salicylate mounting media. The coverslips of the slides were then sealed with nail polish. Channels for the three different wavelengths were set for maximum absorption and were 405 nm for DAPI, 488 nm for GFP, and 561 nm for phalloidin using a Carl Zeiss LSM 710 confocal microscope (Oberkochen, Germany). Voltage gains were set at the same level for each image using ZEN blue software v. 3.5.

### 7. Statistical analyses

All statistical analyses were carried out using JMP Pro v. 18 software and visualizations with GraphPad Prism v.10. The Gustatory Response Scores (GRS) were considered as count data and each day was analyzed with a 2 x 3 chi-square goodness of fit test where responses and non-responses were considered for each treatment group. This was then followed up with *post hoc* 2 x 2 Fisher’s exact tests for each of the pairwise comparisons within each day. The *in vivo* and *in vitro* normalized gene expression data within each treatment were considered to be non-normal, so a Kruskal-Wallis Test was carried out comparing the normalized gene expression levels within each day, across treatments, that was then followed by a post hoc Wilcoxon Rank Sum test for pairwise comparisons.

## Acknowledgements

We would like to thank Dr. Batu Erman and Dr. Emrah Eroğlu for helpful advice and discussions regarding the design of the CRISPR-Cas9 plasmid, as well as Emre Can Gunaydin, Ilknur Bahcivan, Can Sontepe, Faruk Barbaros Kalyoncu, Gizem Sahin, Zeynep Buse Orhan for help with honey bee brain injections and sample processing. We thank Daniel Fisher from Agilent Technologies for his assistance in designing the guide RNAs in the preliminary validation of the use of the CRISPR-Cas9 system in honey bees. The statements made in the article represent the authors’ views and should not be interpreted as endorsement from their respective employers or the funding agencies. The mention of trade names or commercial products in this publication is solely for the purpose of providing specific information and does not imply recommendation or endorsement by the US Department of Agriculture (USDA). USDA is an equal opportunity provider, employer, and lender.

## Ethics

No special ethical approval is required to carry out this research.

## Data Availability

The data collected for this study can be found publicly available with the following DOI 10.17605/OSF.IO/VA7RF.

## Authors Contributions

The original conceptualization of the study was performed by BEKD, CM, and SG. ACS, BEKD, DT, FN, MG, RLB were responsible for carrying out the study and collecting the data. BEKD, CM, FN, and MG conducted the data analysis. CM, MG, and RLB provided the supervision for the study, the research facilities, and software. Funding for the study was obtained by BEKD, CM, and MG. CM and BEKD were primarily responsible for writing the manuscript. All authors contributed to reviewing and editing the manuscript.

## References

1. Adli, M. (2018). The CRISPR tool kit for genome editing and beyond. Nature Communications 9, 1911. 10.1038/s41467-018-04252-2.

2. Lu, L., Shen, X., Sun, X., Yan, Y., Wang, J., and Yuan, Q. (2022). CRISPR-based metabolic engineering in non-model microorganisms. Current Opinion in Biotechnology 75, 102698. 10.1016/j.copbio.2022.102698.

3. Pacesa, M., Pelea, O., and Jinek, M. (2024). Past, present, and future of CRISPR genome editing technologies. Cell 187, 1076–1100. 10.1016/j.cell.2024.01.042.

4. Zou, Y., Sun, X., Yang, Q., Zheng, M., Shimoni, O., Ruan, W., Wang, Y., Zhang, D., Yin, J., Huang, X., et al. (2022). Blood-brain barrier-penetrating single CRISPR-Cas9 nanocapsules for effective and safe glioblastoma gene therapy. Sci Adv 8, eabm8011. 10.1126/sciadv.abm8011.

5. Rasul, M.F., Hussen, B.M., Salihi, A., Ismael, B.S., Jalal, P.J., Zanichelli, A., Jamali, E., Baniahmad, A., Ghafouri-Fard, S., Basiri, A., and Taheri, M. (2022). Strategies to overcome the main challenges of the use of CRISPR/Cas9 as a replacement for cancer therapy. Mol Cancer 21, 64. 10.1186/s12943-021-01487-4.

6. Dai, W.-J., Zhu, L.-Y., Yan, Z.-Y., Xu, Y., Wang, Q.-L., and Lu, X.-J. (2016). CRISPR-Cas9 for *in vivo* Gene Therapy: Promise and Hurdles. Molecular Therapy - Nucleic Acids 5. 10.1038/mtna.2016.58.

7. Leboulle, G., Gehne, N., Froese, A., and Menzel, R. (2022). *In-vivo* egfp expression in the honeybee *Apis mellifera* induced by electroporation and viral expression vector. PLOS ONE 17, e0263908. 10.1371/journal.pone.0263908.

8. Balfanz, S., Jordan, N., Langenstuck, T., Breuer, J., Bergmeier, V., and Baumann, A. (2014). Molecular, pharmacological, and signaling properties of octopamine receptors from honeybee (*Apis mellifera*) brain. J. Neurochem. 129, 284–296. 10.1111/jnc.12619.

9. Xu, J., Xu, X., Zhan, S., and Huang, Y. (2019). Genome editing in insects: current status and challenges. National Science Review 6, 399–401. 10.1093/nsr/nwz008.

10. Chen, Z., Traniello, I.M., Rana, S., Cash-Ahmed, A.C., Sankey, A.L., Yang, C., and Robinson, G.E. (2021). Neurodevelopmental and transcriptomic effects of CRISPR/Cas9-induced somatic *orco* mutation in honey bees. Journal of Neurogenetics 35, 320–332. 10.1080/01677063.2021.1887173.

11. Brahma, A., Frank, D.D., Pastor, P.D.H., Piekarski, P.K., Wang, W., Luo, J.-D., Carroll, T.S., and Kronauer, D.J.C. (2023). Transcriptional and post-transcriptional control of odorant receptor choice in ants. Current Biology 33, 5456–5466.e5455. 10.1016/j.cub.2023.11.025.

12. Kohno, H., Suenami, S., Takeuchi, H., Sasaki, T., and Kubo, T. (2016). Production of Knockout Mutants by CRISPR/Cas9 in the European Honeybee, *Apis mellifera* L. Zoological Science 33, 505. 10.2108/zs160043.

13. Cullen, G., Gilligan, J.B., Guhlin, J.G., and Dearden, P.K. (2023). Germline progenitors and oocyte production in the honeybee queen ovary. Genetics 225. 10.1093/genetics/iyad138.

14. Menzel, R., Leboulle, G., and Eisenhardt, D. (2006). Small Brains, Bright Minds. Cell 124, 237–239.

15. Gould, J.L. (1986). The locale map of honey bees: do insects have cognitive maps? Science 232, 861–863. 10.1126/science.232.4752.861.

16. Paoli, M., Wystrach, A., Ronsin, B., and Giurfa, M. (2024). Smell and Aftersmell: Fast Calcium Imaging Dynamics of Honey Bee Olfactory Coding. eLife Sciences Publications, Ltd.

17. Dyer, A.G., Paulk, A.C., and Reser, D.H. (2011). Colour processing in complex environments: insights from the visual system of bees. Proceedings of the Royal Society B: Biological Sciences 278, 952–959. 10.1098/rspb.2010.2412.

18. Chittka, L. (2017). Bee cognition. Curr. Biol. 27, R1049–R1053. 10.1016/j.cub.2017.08.008.

19. Menzel, R., and Muller, U. (1996). Learning and Memory in Honeybees: From Behavior to Neural Substrates. Annual Review of Neuroscience 19, 379–404.

20. Airenne, K.J., Hu, Y.-C., Kost, T.A., Smith, R.H., Kotin, R.M., Ono, C., Matsuura, Y., Wang, S., and Ylä-Herttuala, S. (2013). Baculovirus: an Insect-derived Vector for Diverse Gene Transfer Applications. Molecular Therapy 21, 739–749. 10.1038/mt.2012.286.

21. Chen, C.Y., Lin, C.Y., Chen, G.Y., and Hu, Y.C. (2011). Baculovirus as a gene delivery vector: recent understandings of molecular alterations in transduced cells and latest applications. Biotechnol Adv 29, 618–631. 10.1016/j.biotechadv.2011.04.004.

22. Schulte, C., Leboulle, G., Otte, M., Grünewald, B., Gehne, N., and Beye, M. (2013). Honey bee promoter sequences for targeted gene expression. Insect Mol Biol 22, 399–410. 10.1111/imb.12031.

23. Xu, J., Xu, X., Zhan, S., and Huang, Y. (2019). Genome editing in insects: current status and challenges. Natl Sci Rev 6, 399–401. 10.1093/nsr/nwz008.

24. Bai, Y., He, Y., Shen, C.Z., Li, K., Li, D.L., and He, Z.Q. (2023). CRISPR/Cas9-Mediated genomic knock out of tyrosine hydroxylase and yellow genes in cricket Gryllus bimaculatus. PLoS One 18, e0284124. 10.1371/journal.pone.0284124.

25. Li, J.-J., Shi, Y., Wu, J.-N., Li, H., Smagghe, G., and Liu, T.-X. (2021). CRISPR/Cas9 in lepidopteran insects: Progress, application and prospects. J Insect Physiol 135, 104325. 10.1016/j.jinsphys.2021.104325.

26. Guo, X., Wang, Y., Sinakevitch, I., Lei, H., and Smith, B.H. (2018). Comparison of RNAi knockdown effect of tyramine receptor 1 induced by dsRNA and siRNA in brains of the honey bee, Apis mellifera. J Insect Physiol 111, 47–52. 10.1016/j.jinsphys.2018.10.005.

27. Quigley, T., and Amdam, G. (2024). Ultrastructural Organization of the Honeybee Blood-Brain Barrier and Comparison with Age. bioRxiv, 2024.2009.2027.615080. 10.1101/2024.09.27.615080.

28. Ikeda, T., Nakamura, J., Furukawa, S., Chantawannakul, P., Sasaki, M., and Sasaki, T. (2011). Transduction of baculovirus vectors to queen honeybees, Apis mellifera. Apidologie 42, 461–471. 10.1007/s13592-011-0014-z.

29. Akülkü, İ. (2021). Mapping the distribution of the octopamine beta receptor subtype-2 in the honeybee (apis mellifera) brain.

30. Muenz, T.S., Groh, C., Maisonnasse, A., Le Conte, Y., Plettner, E., and Rössler, W. (2015). Neuronal plasticity in the mushroom body calyx during adult maturation in the honeybee and possible pheromonal influences. Dev Neurobiol 75, 1368–1384. 10.1002/dneu.22290.

31. Naik, N.G., Lo, Y.-W., Wu, T.-Y., Lin, C.-C., Kuo, S.-C., and Chao, Y.-C. (2018). Baculovirus as an efficient vector for gene delivery into mosquitoes. Scientific Reports 8, 17778. 10.1038/s41598-018-35463-8.

32. Bougioukli, S., Chateau, M., Morales, H., Vakhshori, V., Sugiyama, O., Oakes, D., Longjohn, D., Cannon, P., and Lieberman, J.R. (2021). Limited potential of AAV-mediated gene therapy in transducing human mesenchymal stem cells for bone repair applications. Gene Therapy 28, 729–739. 10.1038/s41434-020-0182-4.

33. Kalesnykas, G., Kokki, E., Alasaarela, L., Lesch, H.P., Tuulos, T., Kinnunen, K., Uusitalo, H., Airenne, K., and Yla-Herttuala, S. (2017). Comparative Study of Adeno-associated Virus, Adenovirus, Bacu lovirus and Lentivirus Vectors for Gene Therapy of the Eyes. Curr Gene Ther 17, 235–247. 10.2174/1566523217666171003170348.

34. Ando, T., Fujiyuki, T., Kawashima, T., Morioka, M., Kubo, T., and Fujiwara, H. (2007). In vivo gene transfer into the honeybee using a nucleopolyhedrovirus vector. Biochem Biophys Res Commun 352, 335–340. 10.1016/j.bbrc.2006.11.020.

35. Robinson, K.O., Ferguson, H.J., Cobey, S., Vaessin, H., and Smith, B.H. (2000). Sperm-mediated transformation of the honey bee, *Apis mellifera*. Insect Mol Biol 9, 625–634. 10.1046/j.1365-2583.2000.00225.x.

36. Kohno, H., Suenami, S., Takeuchi, H., Sasaki, T., and Kubo, T. (2016). Production of Knockout Mutants by CRISPR/Cas9 in the European Honeybee, *Apis mellifera* L. Zoological science 33, 505–512. 10.2108/zs160043.

37. Sprink, T., and Wilhelm, R. (2024). Genome Editing in Biotech Regulations Worldwide. In A Roadmap for Plant Genome Editing, A. Ricroch, D. Eriksson, D. Miladinović, J. Sweet, K. Van Laere, and E. Woźniak-Gientka, eds. (Springer Nature Switzerland), pp. 425-435. 10.1007/978-3-031-46150-7_25.

38. Chen, Z., Traniello, I.M., Rana, S., Cash-Ahmed, A.C., Sankey, A.L., Yang, C., and Robinson, G.E. (2021). Neurodevelopmental and transcriptomic effects of CRISPR/Cas9- induced somatic orco mutation in honey bees. J Neurogenet 35, 320–332. 10.1080/01677063.2021.1887173.

39. Mayack, C., Natsopoulou, M.E., and McMahon, D.P. (2015). *Nosema ceranae* alters a highly conserved hormonal stress pathway in honeybees. Insect Molecular Biology 24, 662–670. 10.1111/imb.12190.

40. Kunieda, T., and Kubo, T. (2004). In vivo gene transfer into the adult honeybee brain by using electroporation. Biochemical and Biophysical Research Communications 318, 25–31. 10.1016/j.bbrc.2004.03.178.

41. Akülkü, İ., Ghanem, S., Filiztekin, E., Suwannapong, G., and Mayack, C. (2021). Age- Dependent Honey Bee Appetite Regulation Is Mediated by Trehalose and Octopamine Baseline Levels. Insects 12. 10.3390/insects12100863.

42. Mayack, C., Phalen, N., Carmichael, K., White, H.K., Hirche, F., Wang, Y., Stangl, G.I., and Amdam, G.V. (2019). Appetite is correlated with octopamine and hemolymph sugar levels in forager honeybees. Journal of comparative physiology. A, Neuroethology, sensory, neural, and behavioral physiology 205, 609–617. 10.1007/s00359-019-01352-2.

43. Ghanem, S., Akülkü, İ., Güzle, K., Khan, Z., and Mayack, C. (2024). Regulation of forager honey bee appetite independent of the glucose-insulin signaling pathway. Frontiers in Insect Science 4. 10.3389/finsc.2024.1335350.

44. Kaya-Zeeb, S., Engelmayer, L., Straßburger, M., Bayer, J., Bähre, H., Seifert, R., Scherf-Clavel, O., and Thamm, M. (2022). Octopamine drives honeybee thermogenesis. eLife 11. 10.7554/eLife.74334.

45. Mansouri, M., and Berger, P. (2018). Baculovirus for gene delivery to mammalian cells: Past, present and future. Plasmid 98, 1–7. 10.1016/j.plasmid.2018.05.002.

46. Ghosh, S., Parvez, M.K., Banerjee, K., Sarin, S.K., and Hasnain, S.E. (2002). Baculovirus as Mammalian Cell Expression Vector for Gene Therapy: An Emerging Strategy. Molecular Therapy 6, 5–11. 10.1006/mthe.2000.0643.

47. Chen, L., Wang, G., Zhu, Y.N., Xiang, H., and Wang, W. (2016). Advances and perspectives in the application of CRISPR/Cas9 in insects. Dongwuxue Yanjiu 37, 220–228. 10.13918/j.issn.2095-8137.2016.4.220.

48. Dow, L.E., Fisher, J., O’Rourke, K.P., Muley, A., Kastenhuber, E.R., Livshits, G., Tschaharganeh, D.F., Socci, N.D., and Lowe, S.W. (2015). Inducible in vivo genome editing with CRISPR-Cas9. Nat. Biotechnol. 33, 390–394. 10.1038/nbt.3155.

49. Mayack, C., and Naug, D. (2011). A changing but not an absolute energy budget dictates risk-sensitive behaviour in the honeybee. Animal Behaviour 82, 595–600.

50. Søvik, E., Plath, J.A., Devaud, J.M., and Barron, A.B. (2016). Neuropharmacological Manipulation of Restrained and Free-flying Honey Bees, Apis mellifera. Journal of visualized experiments : JoVE. 10.3791/54695.

51. Bitterman, M.E., Menzel, R., Fietz, A., and Schäfer, S. (1983). Classical conditioning of proboscis extension in honeybees (*Apis mellifera*). J. Comp. Psychol. 97, 107–119.

52. Lourenço, A.P., Mackert, A., Cristino, A.D., and Simoes, Z.L.P. (2008). Validation of reference genes for gene expression studies in the honey bee, *Apis mellifera*, by quantitative real-time RT-PCR. Apidologie 39, 372–U333. 10.1051/apido:2008015.

