## Supplemental File for "Successful delivery of CRISPR-Cas9 with a baculovirus vector for insect brain targets"

**Supplemental Information**

1. *Preliminary testing of CRISPR-Cas9 on the octopamine beta receptor subtypes*
   1.1 *Material and Methods*

*1.2 Guide RNA design*

Three single guide RNAs (sgRNAs) were designed for each of the four *octopamine beta receptor subtypes* found in honey bees (*AmOctB1-4R*), yielding twelve guides in total (**Table S1**). Three sgRNAs per target was chosen to increase the probability of causing a significant mutation in the target gene. All guides were used at once to maximize the probability of a successful editing. The guides consisted of two main sections. The first was a 20-nucleotide base long sequence that aligned with the target sequence. Each guide was then designed to align with a portion of the genome that was followed by a PAM region, encoded by “NGG” nucleotides where N can be any nucleotide base. Following the portion of the sgRNA that anneals to the target sequence there was an eighty-base conserved sequence that corresponded to a piece of the Cas9 enzyme. This portion of the sgRNA anchors to the Cas9 enzyme, directing it to the desired cut site. The final three nucleotide bases of each sgRNA sequence were modified with an MS motif. This modification involved replacing the hydrogen at the second position on the sugar portion of the backbone, with a methoxy group, and replacing one negatively charged oxygen in the phosphate portion of the backbone with a negatively charged sulfur. These modifications confer stability to the sgRNAs and work to strengthen their hold on the Cas9 enzyme when complexed. The sgRNAs were designed and ordered from Agilent Technologies (Santa Clara, CA, USA).

**Table S1. s**gRNA designs in which each guide had a portion that matched with the target sequence as well as a portion that anchors the guide to the enzyme. * indicates that the base was stabilized by an MS motif.

| **Target Gene** | **sgRNA sequence 5’ -> 3’** | **Target sequence 5” -> 3’** |
| --- | --- | --- |
| *AmOctβ1R* | GGAAGCGAACGAAACGACGGGUUUUAGAGCUAGAAAUA GCAAGUUAAAAUAAGGCUAGUCCGUUAUCAACUUGAAA AAGUGGCACCGAGUCGGUGCUU*U*U*  CCGCGAGGAGAACGACGAAUGUUUUAGAGCUAGAAAUA GCAAGUUAAAAUAAGGCUAGUCCGUUAUCAACUUGAAA  AAGUGGCACCGAGUCGGUGCUU*U*U*  CGAAAAGCUGCGAGAACGUUGUUUUAGAGCUAGAAAUA GCAAGUUAAAAUAAGGCUAGUCCGUUAUCAACUUGAAA  AAGUGGCACCGAGUCGGUGCUU*U*U* | GGAAGCGAACGAAACG ACGGAGG  CCGCGAGGAGAACGACG AATTGG  CGAAAAGCTGCGAGAAC GTTAGG |
| *AmOctβ2R* | GACGAGCAGCGAAUCGAGCGGUUUUAGAGCUAGAAAUA GCAAGUUAAAAUAAGGCUAGUCCGUUAUCAACUUGAAA AAGUGGCACCGAGUCGGUGCUU*U*U*  GACGUGACGACCCUGUUGAAGUUUUAGAGCUAGAAAUA GCAAGUUAAAAUAAGGCUAGUCCGUUAUCAACUUGAAA  AAGUGGCACCGAGUCGGUGCUU*U*U*  GAACGGGGAGUUGAACAGCGGUUUUAGAGCUAGAAAUA GCAAGUUAAAAUAAGGCUAGUCCGUUAUCAACUUGAAA  AAGUGGCACCGAGUCGGUGCUU*U*U* | GACGAGCAGCGAATCGA GCGAGG  GACGTGACGACCCTGTT GAACGG  GAACGGGGAGTTGAACA GCGCGG |
| *AmOctB3R* | AGUCAGUGAGCCAUCGGACGGUUUUAGAGCUAGAAAUA GCAAGUUAAAAUAAGGCUAGUCCGUUAUCAACUUGAAA AAGUGGCACCGAGUCGGUGCUU*U*U*  ACCGAUCAACGUCGUUUGGAGUUUUAGAGCUAGAAAUA GCAAGUUAAAAUAAGGCUAGUCCGUUAUCAACUUGAAA AAGUGGCACCGAGUCGGUGCUU*U*U*  GAGGACGCUCGGAAUAAUAAGUUUUAGAGCUAGAAAUA  GCAAGUUAAAAUAAGGCUAGUCCGUUAUCAACUUGAAA AAGUGGCACCGAGUCGGUGCUU*U*U* | AGTCAGTGAGCCATCGG ACGAGG  ACCGATCAACGTCGTTT GGACGG  GAGGACGCTCGGAATAA TAATGG |
| *AmOctβ4R* | GACGAGUCUGCCCAGCCGAGGUUUUAGAGCUAGAAAUA GCAAGUUAAAAUAAGGCUAGUCCGUUAUCAACUUGAAA AAGUGGCACCGAGUCGGUGCUU*U*U*  CGCUGCGAACAACGUCACCUGUUUUAGAGCUAGAAAUA GCAAGUUAAAAUAAGGCUAGUCCGUUAUCAACUUGAAA AAGUGGCACCGAGUCGGUGCUU*U*U*  GAGCCGAGCACAAAGCUGCGGUUUUAGAGCUAGAAAUA GCAAGUUAAAAUAAGGCUAGUCCGUUAUCAACUUGAAA  AAGUGGCACCGAGUCGGUGCUU*U*U* | GACGAGTCTGCCCAGCC GAGTGG  CGCTGCGAACAACGTCA CCTCGG  GAGCCGAGCACAAAGCT GCGA |

*1.3 Honey bee sampling*

In-hive honey bees were collected from a brood frame using a 50-mL conical tube, from three source colonies, from two different apiaries, located in Oxford (40.0379° N, 76.3055° W) and The Grayson School Apiary in Radnor (40.0462° N, 75.3599° W), Pennsylvania. The source colonies were considered to be in good health due to the low numbers of mites, high volume of eggs and brood, and presence of a queen. Honey bees were transferred into small metal cages and were given a sucrose solution as a food source. The honey bees were then brought back to Haverford College and stored in an incubator set at 37 ºC and 60% Relative Humidity until experiments began the following day.

*1.3 CRISPR-Cas9 and guide RNA injections*

Honey bees were removed from the incubator and placed into a 20 mL glass scintillation vial onto crushed ice until immobilized. Individual honey bees were then harnessed into conventional, store-bought, drinking straws that were cut to ~3 inches in length. The necks of the honey bee were strapped to the straw using 2-3 mm thin, 70 mm long strips of Gorilla Duct Tape. Once harnessed, the bees were given a 30 min acclimation period and then they were handfed 10 μL of 50% sucrose solution immediately using a micropipetter, as well as every 12 h until they were euthanized. A total of 63 honey bees at one time were harnessed to carry out the experiment.

1.4 *CRISPR-Cas9 and guide RNA injections*

sgRNAs were reconstituted with TE buffer (10 mM Tris, 1 mM EDTA, pH 8) upon arrival. sgRNAs were diluted to 50 μM stock solutions at a pH of 7.4. Solutions were held at room temperature to dissolve the guides and then vortexed. sgRNAs were then flash frozen with liquid nitrogen and stored at -80 ºC. In a 1.5 mL microcentrifuge tube, 2.5 μg Cas9 nuclease (Aldevron, Fargo, ND, USA) and 500 ng of each sgRNA (Agilent Technologies, Santa Clara, CA, USA), was dissolved in 50 μL of Opti-Mem I Medium (Thermo Fisher Scientific, Waltham, MA, USA). Then 5 μL of Cas9 Plus was added to the same tube and vortexed. All twelve sgRNAs were added in concert. In another 1.5 mL microcentrifuge tube, 3 μL of Lipofectamine CRISPRMAX (Thermo Fisher Scientific, Waltham, MA, USA) was diluted in 50 μL of Opti-Mem I Medium (Thermo Fisher Scientific, Waltham, MA, USA). This tube was then immediately added to the other, and was incubated at room temperature for 10 min. A total of 100 μL of this CRISPR-Cas9 solution was loaded into a 10 μL Hamilton syringe outfitted with a 30-gauge needle (Thermo Fisher Scientific, Waltham, MA, USA). Each harnessed honey bee received a 1 μL injection except for the sham control which received a 1 μL injection of TE buffer. A third of the honey bees received an ocellar tract injection into their head (N = 21). The other third received injections to their thorax (N = 21), where the solution could be taken up into the open circulatory system. The last third received the sham injection into the thorax (N = 21). After each honey bee was injected they were fed 10 μL of 50% sucrose solution every 12 hours and returned to the incubator set at 37 ºC and 60% Relative Humidity. At 8, 24, and 48 h, after injection, half of the remaining honey bees were euthanized with liquid nitrogen and then transferred to a -80 ºC freezer for long term storage until further analysis.

1.5 *PCR amplification and Sanger sequencing of the AMOctβ1-4R target genes*

From each whole bee that was injected, the DNA was extracted using a Qiagen (Hilden, Germany) DNeasy kit following the manufacturer instructions. For the PCR reactions, one set of primers per *AMOctβ1-4* gene target was self-designed and validated with a previous qPCR melt curve analysis (**Table S2**). The following PCR reaction was carried out for each gene target: 2 μL of forward and reverse primers (10 mM) (Thermo Fisher Scientific, Waltham, MA, USA), 25 μL of PCR 2x MasterMix (Thermo Scientific, Waltham, MA, USA), 9 μL of nuclease free water, and 10 μL of template DNA was combined for a 50 uL final volume reaction. These reactions were carried out in triplicate with negative controls. The Biorad T100 thermocycler (Hercules, CA, USA) was programmed with the following parameters: 3 min at 95 ºC denaturization step, 40 cycles of 30 s at 95 ºC, 0:30 s at 55 ºC, and 30 s at 72 ºC, followed by 30 s at 95 ºC and 1 min at 60 ºC to elongate amplified strands. The samples were then held at 4 ºC indefinitely. The amplified targets were then isolated using gel electrophoresis with a 2% agarose gel and TBE buffer (89 mM Tris, 89 mM boric acid, 2 mM EDTA). A total of 10 μL of each sampled was loaded into each well and were then run for 45 min. An Ultra Low Range DNA Ladder (Thermo Scientific, Waltham, MA, USA) was used as a reference for estimating the size of bands. The remaining 40 μL of the samples were purified using a PCR Purification Kit (Qiagen, Hilden, Germany) following the manufacturer instructions. From the PCR-purified products, 50 μL was sent to Genewiz for Sanger sequencing. The returned sequences were subjected to a NCBI’s BLAST sequence alignment to determine if amplified sequences matched the targets. The results of sequencing are shown in **Table S3**.

**Table S2**. Primer sequences with corresponding melt temperatures and amplicon lengths.

| **Gene** | **Forward Primer** | **Reverse Primer** | **Tm** | **Amplicon length (bp)** |
| --- | --- | --- | --- | --- |
| *AmOctß1R*  HF548209.1 | TTCTGGGTGGGGTACTTCAG | CCTGCTTCCTATGAGCTTGG | 60 | 151 |
| *AmOctß2R*  HF548210.1 | ACAATTTGAACGGGGAGTTG | AAGAAGGGCAACCAGCATAG | 60 | 187 |
| *AmOctß3R*  HF548211.1 | ATCCAGCGATCATGAAGAGG | TCACCTCGAACGTGCATATC | 60 | 162 |
| *AmOctß4R*  HF548212.1 | TGAGAGTGGTGACGAATTGC | GAATGGAGGCTGTGGAAAAG | 60 | 177 |

2. *Results*

All four octopamine beta receptor subtypes (*AmOctβ1-4R*) had successful mutations caused from the CRISPR-Cas9 injections from at least one guide RNA, whether this was a point mutation or an indel modification. An indel mutation is likely to cause the protein to be non-functional. A representation of each mutation can be found in Table S3.

**Table S3.** The Sanger sequencing results obtained from Genewiz which specifies where the honey bee was injected with the CRISPR-Cas9 system, the gene target it is intended for, which of the three guide RNAs that was used, the alleles that were modified, the alteration of the sequence highlighted in red, and whether the sequencing product could not be produced due to a indel mutation is provided in the comments column.

| **Injection site** | **Gene target** | **sgRNA** | **Alleles** | **Flanking sequence** | **Comments** |
| --- | --- | --- | --- | --- | --- |
| Head | AmOctβ1R | 1 | C/C | CGACGGAGGATGGATTGGTGG |  |
| Thorax | AmOctβ1R | 1 |  |  | Cannot call due to indel |
| Head | AmOctβ1R | 2 | C/C | TGGAGCGCGTTATTCTTCTGA |  |
| Thorax | AmOctβ1R | 2 | C/C | TGGAGCGCGTTATTCTTCTGA | Called off of R trace due to indel |
| Head | AmOctβ1R | 3 | AC/- | ACTGTTATATACATATATAAAT |  |
| Thorax | AmOctβ1R | 3 | AC/- | ACTGTTATATACATATATAAAT |  |
| Head | AmOctβ2R | 1 | -/- | CTAAGGAGCAGAGGAGGAGGAGG |  |
| Thorax | AmOctβ2R | 1 | -/- | CTAAGGAGCAGAGGAGGAGGAGG |  |
| Head | AmOctβ2R | 2 |  |  | No mutations observed |
| Thorax | AmOctβ2R | 2 |  |  | No mutations observed |
| Head | AmOctβ2R | 3 | T/T | AGGATTTGAACAATTTGAACG |  |
| Thorax | AmOctβ2R | 3 | T/T | AGGATTTGAACAATTTGAACG |  |
| Head | AmOctβ3R | 1 |  |  | No mutations observed |
| Thorax | AmOctβ3R | 1 |  |  | No mutations observed |
| Head | AmOctβ3R | 2 | G/G | CGTTGGACGTCTACTTCTCCA |  |
| Thorax | AmOctβ3R | 2 | T/T | ACGTCTACTTCTCCACGGCCT |  |
| Head | AmOctβ3R | 3 |  |  | No mutations observed |
| Thorax | AmOctβ3R | 3 |  |  | No mutations observed |
| Head | AmOctβ4R | 1 |  |  | No mutations observed |
| Thorax | AmOctβ4R | 1 |  |  | No mutations observed |
| Head | AmOctβ4R | 2 | G/G | TGTTCGTGGTAGTGATCAAAG |  |
| Thorax | AmOctβ4R | 2 | A/G | TGTTCGTGGTAGTGATCAAAG |  |
| Head | AmOctβ4R | 3 |  |  | No mutations observed |
| Thorax | AmOctβ4R | 3 |  |  | No mutations observed |

2 *Baculovirus Plasmid Design*

The CRISPR-Cas9 system targeting the *AmOctB2* receptor was integrated in the baculovirus vector (VB210408-1080vfc) (Figure S1). The system was integrated into the baculovirus *Autographa californica nucleopolyhedrovirus* (AcNPV) genome by the VectorBuilder company.

CRISPRflydesign was used to optimize the gRNA with regards to off-target and on-target sites. Because the honey bee (*Apis mellifera*) was not available on CRISPRflydesign, nor on others such as CHOP-CHOP, Benchling was used to design the gRNA instead. The beta-octopamine subtype 2 receptor sequence was uploaded to Benchling. Preference was given to gRNAs located at the start of the octopamine receptor sequence. This approach was followed to avoid the potential of creating a truncated protein. The gRNA promotor of choice was dU6-3. This was the most expressed of the three U6 promoters that have been optimized for the fruit fly (Huynh, Depner, Larson, & King-Jones, 2020).

To ensure high expression levels, each element requires its own promoter, which adds complexity to the plasmid design. This challenge was addressed by integrating a T2A sequence. This allows for the translation of multicistronic sequences, enabling multiple genes to be expressed from a single promoter. T2A causes the ribosomes to skip synthesis of the peptide bond that would normally (based on proposed plasmid design) connect the two proteins, Cas9 with the EGFP-PAC fusion protein, resulting in the synthesis of two separate proteins without the use of multiple promoters.

The *Drosophila melanogaster* Cas9 is highly conserved compared to AmCas9 and has been demonstrated to be effective in past studies. The Tn7l and Tn7r elements allow for the integration of this plasmid into the baculovirus. PucOri replaced the Ori sequence that was originally included along with the addition of Gentamicin. These elements were essential for the VectorBuilder company (Chicago, IL, USA) to synthesize the plasmid and to ensure the successful knock down of the beta octopamine subtype 2 receptor.

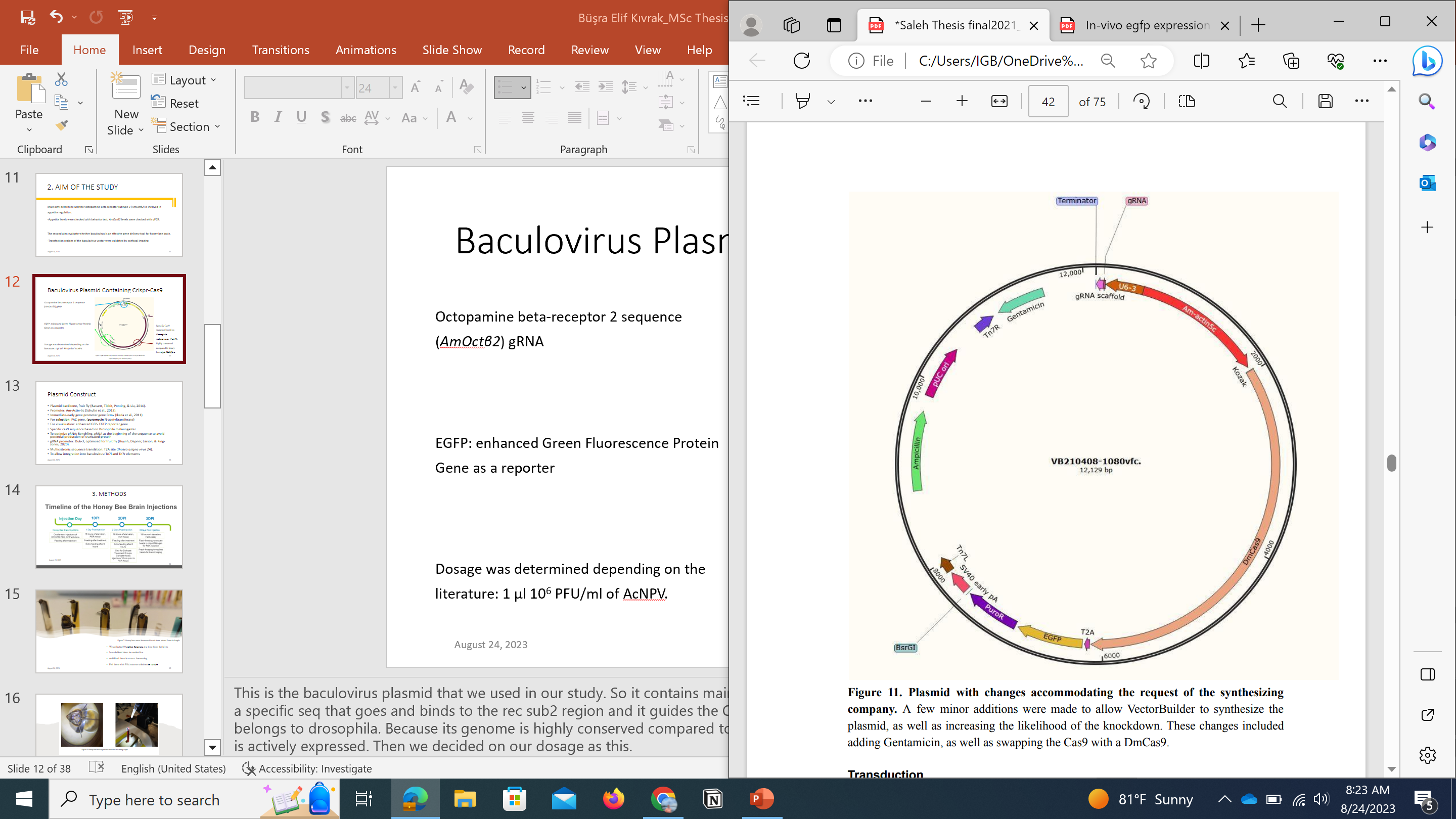

**Figure S1.** **pAC-sgRNA-Cas9 plasmid containing CRISPR system to target *AmOctβ2R*.** The CRISPR-Cas9 system was designed to knockdown *AmOctβ2R*. The system was integrated into the baculovirus *Autographa californica nucleopolyhedrovirus* (AcNPV) genome. The AcNPV vector was prepared by the VectorBuilder company. They provided a specific Cas9 sequence based on *Drosophila melanogaster* as its genome is highly conserved compared to *Apis Mellifera*. Guide RNA for the CRISPR system was prepared with the Benchling program by selecting *Apis mellifera* as the target genome. The Octopamine beta-receptor 2 sequence needed for the gRNA design was obtained from literature^1^. EGFP sequence was also inserted into the system as the reporter of gene editing.

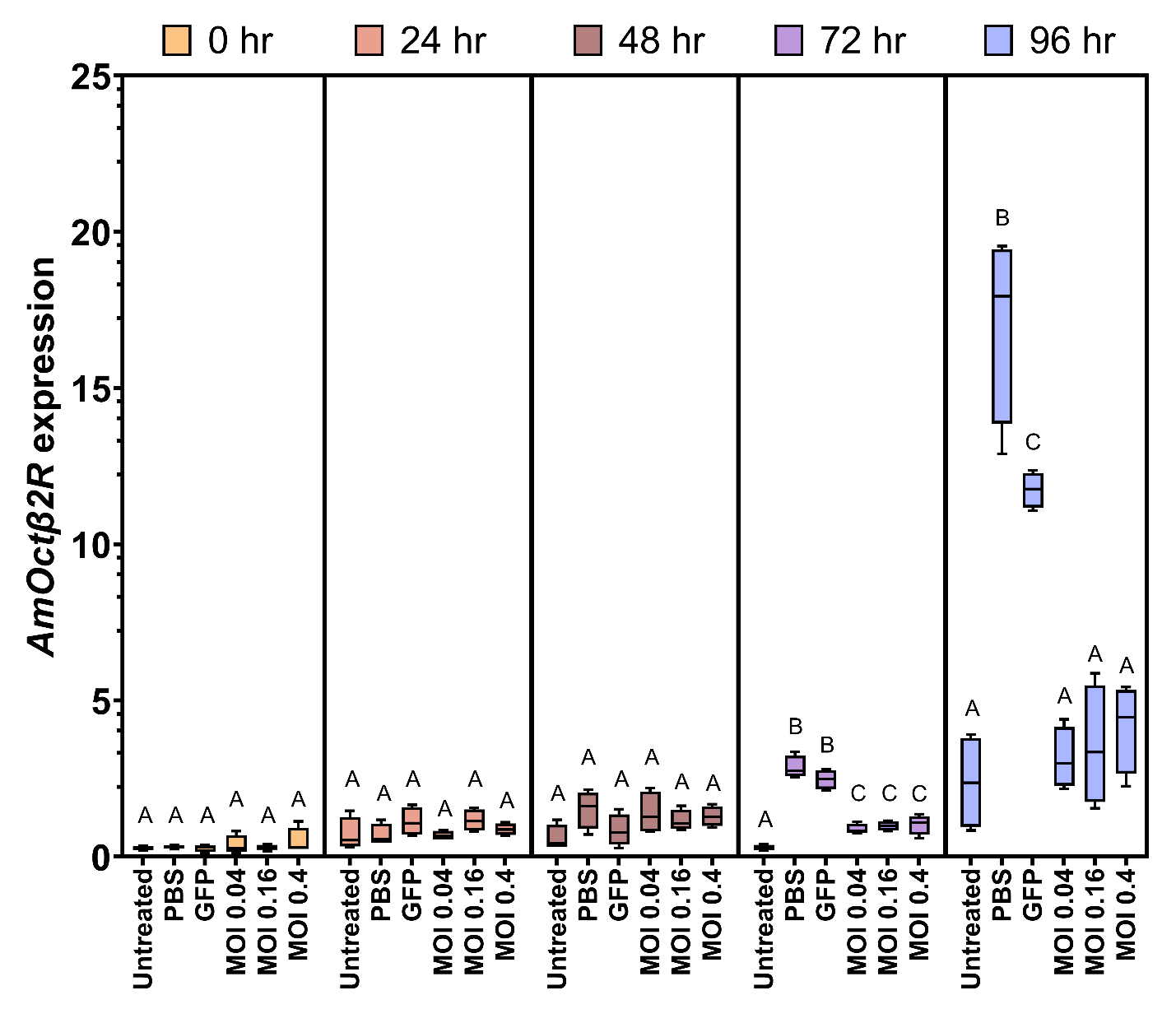

**Figure S2.** Normalized expression of octopamine beta receptor subtype 2 (*AmOctβ2R*) from the honey bee (*Apis mellifera*) cell line, AmE-711, from 0 to 96 hours after treatment in 24-well microplates. Untreated controls received no treatment except medium change only, while 1x phosphate buffer solution (PBS) controls, CRISPR-cas9 green fluorescent protein (GFP) controls, CRISPR-Cas9 *AmOctβ2R* knock down treated bees with multiplicity of infection (MOI) of 0.04, 0.16, and 0.4 baculovirus vector concentrations were treated at the start of the experiment (0 hr). Boxes represent the interquartile range and the middle horizontal line represents the median; whiskers represent the minimum and maximum values. The letters above the boxes represent significant differences at the α = 0.05 level. Each box represents four replicates within each treatment per time point.

**Table S1.** Primer pairs used in the qPCR assay to measure the *in vivo* and *in vitro* knockdown of *AmOctβ2R* normalized to the reference genes *RP49* and *Rps5*, respectively.

| Gene | Sequence (Forward) | Sequence (Reverse) | Reference |
| --- | --- | --- | --- |
| *RP49* | 5’CGTCATATGTTGCCAACTGGT 3’ | 5’TTGAGCACGTTGAACAATGG 3’ | ^2^ |
| *AmOctβ2R* | 5’ACAATTTGAACGGGGAGTTG 3’ | 5’AAGAAGGGCAACCAGCATAG 3’ | Self-designed |
| *Rps5* | 5’AATTATTTGGTCGCTGGAATTG 3’ | 5’TAACGTCCAGCAGAATGTGGTA 3’ | ^3^ |

**References**

1.	Balfanz, S., Jordan, N., Langenstuck, T., Breuer, J., Bergmeier, V., and Baumann, A. (2014). Molecular, pharmacological, and signaling properties of octopamine receptors from honeybee (*Apis mellifera*) brain. J. Neurochem. *129*, 284-296. 10.1111/jnc.12619.

2.	Lourenço, A.P., Mackert, A., Cristino, A.D., and Simoes, Z.L.P. (2008). Validation of reference genes for gene expression studies in the honey bee, *Apis mellifera*, by quantitative real-time RT-PCR. Apidologie *39*, 372-U333. 10.1051/apido:2008015.

3.	Hamiduzzaman, M.M., Guzman-Novoa, E., and Goodwin, P.H. (2010). A multiplex PCR assay to diagnose and quantify *Nosema* infections in honey bees (*Apis mellifera*). Journal of Invertebrate Pathology *105*, 151-155.
